# On attaining and estimating steady walking speed

**DOI:** 10.64898/2026.03.01.708843

**Authors:** Jeremy D Wong, Osman Darici, Arthur D Kuo

## Abstract

Gait speed is widely used to quantify mobility and functional status, compare how interventions work, and predict health outcomes. Yet despite its ubiquity, the common assumption that short walking tests capture a person’s “steady” speed has rarely been verified. A lack of standardization also makes it difficult to compare gait test results across protocols. Here we show that most gait test distances, typically 10 m or less, are too short for young healthy adults (N = 10) to attain steadiness. Using gait measurements for a range of ten short distances, we found that peak speed increases systematically with total walking distance and only gradually approaches the individual’s preferred speed at about 10 m. How the individual accelerates is characterized by a “distance constant”, also identified from data and only loosely correlated with preferred speed (*ρ* = 0.55). The speed trajectories also show how most conventional gait tests systematically underestimate steady speed by about 30%, while dynamic start and stop conditions partially but not fully corrected the bias. Gait speed is not a single fixed property but a dynamic process that depends on the distance available for acceleration and deceleration. Although conventional gait tests remain valuable, most characterize transient rather than steady walking performance. We further show that measurements at three to five distances are sufficient to predict steady-state speed in young adults with zero average bias. This correction requires only a stopwatch and modest additional testing time, making it readily adoptable in both clinical and research settings.

## Introduction

Walking speed is not only a matter of getting somewhere but also a widely recognized indicator of mobility, functional status, and overall health. Self-selected gait speed predicts a wide range of health outcomes such as recovery after stroke (Schmid et al., 2007; Tilson et al., 2010), mortality with age or medical intervention (Afilalo et al., 2016; White et al., 2013), and incidence of cognitive decline and dementia (Grande et al., 2019). It is a simple yet powerful measure, sometimes called a “sixth vital sign” (Fritz and Lusardi, 2009).

Despite their importance, gait tests are inconsistently defined, with no agreement on when steady speed is attained nor on whether to include transient acceleration. Consequently, measured values often depend as much on the testing protocol as on the individual being tested. Without a clear understanding of what gait tests measure, speeds cannot be compared across studies as objectively as established vital signs such as blood pressure. A quantitative description of both steady speed and how it is attained could make gait assessment more interpretable and standardized.

Gait tests vary widely in distance and start and stop protocols (Fig. 1A). Although speed may be timed over practically any distance, most gait tests are relatively short, with distances of 4 – 10 m most common (Bohannon and Williams Andrews, 2011; Peel et al., 2013; Mehmet et al., 2020). It is generally assumed that steady speed is attained following a brief acceleration interval (Fig. 1B after Breniere and Do, 1986) and prior to deceleration to stop. As such, test protocols may employ a static start explicitly including acceleration, or a dynamic start that excludes some initial distance (Ji et al., 2024; Johnson et al., 2020), dependent on evaluator’s intent. Timing distances may also vary due to constraints at test sites such as for clinics or homes, contributing to twenty unique distances in one survey of forty-one studies (Bohannon and Williams Andrews, 2011). Although the timed distance should be immaterial if gait is indeed steady, the comfortable speeds for adults at least 70 yr spanned a 7.8-fold range in another survey of forty-two studies (Peel et al., 2013). Noting that protocol accounts for much of the differences, authors have repeatedly called for greater standardization (e.g, Fritz and Lusardi, 2009; Graham et al., 2008a; Krumpoch et al., 2021; Peel et al., 2013; Stuck et al., 2020).

**Fig. 1.**
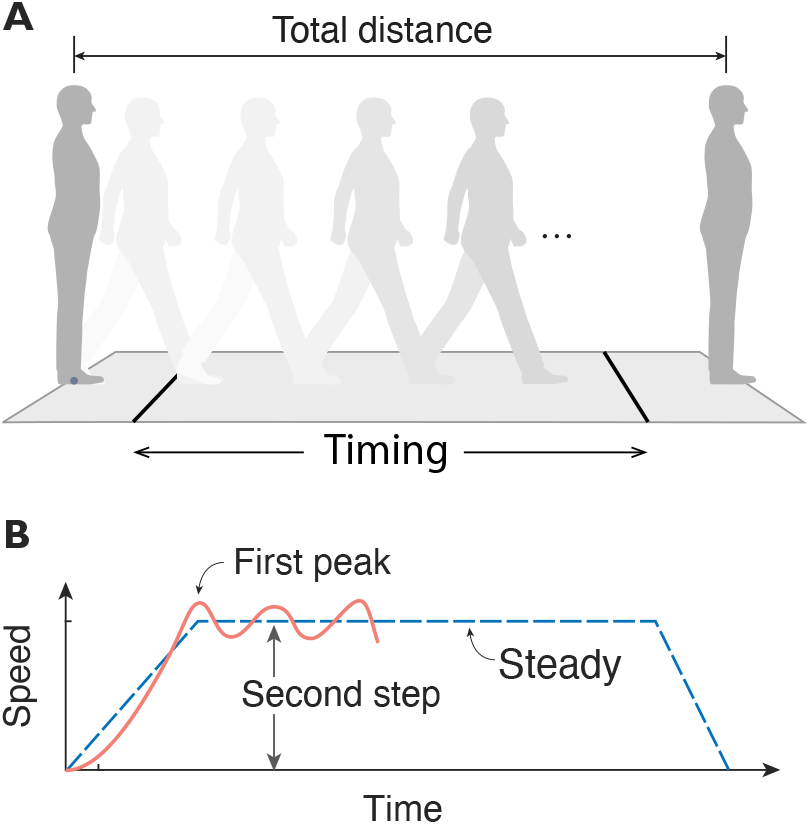
Standard schema for measuring gait speed and steady walking. (A) Speed is typically assessed over a fixed timing distance following gait initiation. Initiation may be defined with a static start from upright rest, or after a dynamic start with an untimed acceleration interval. (B) The speed-time profile is assumed to comprise rapid acceleration, a steady interval, and rapid deceleration (dashed line). Sample data from Breniere & Do (1975) illustrate this profile and two assumed indicators of steady gait: first peak in speed and the average speed of the second step.

Standardization is, however, impeded by a lack of quantitative data. Dynamic start distances are typically specified by convention rather than data, and dynamic stops are often vague or unreported (Graham et al., 2008b; Mehmet et al., 2020; Sustakoski et al., 2015). The prevailing assumption, following Brenere & Do (1986), is that steady speed is achieved within the first step or two, despite that study not directly measuring steady-state walking. Subsequent work suggests otherwise: Lindemann et al. (2008) found that frail older adults required at least 2.5 m to reach steady speed, longer than most current test protocols. Zhao et al. (2021) showed data similar to Breniere & Do (1986) but suggesting longer acceleration intervals. Our own measurements in healthy younger adults suggest that steady speed depends on the total test distance, contrary to conventional assumptions (Fig. 1B). While steadiness may appear obvious on a treadmill or long sidewalk, it has rarely been quantified for short distances typical of gait tests.

Consequently, current gait speed measures cannot be compared objectively across protocols. This study addresses that gap by quantifying how healthy adults attain steady speed within the short distances typical of gait tests. We analyzed full speed trajectories to characterize how gait is initiated and terminated, and how steady speed emerges. From these data, we define an objective *preferred speed* that is independent of test protocol. We further introduce a *distance constant* that describes how rapidly an individual approaches that speed. These measures are used to quantify the bias and variance introduced by conventional gait metrics. Together, these analyses provide a quantitative foundation for defining steady walking and improving the consistency and interpretability of gait speed as a physiological and clinical indicator.

## Methods

We analyzed forward speed trajectories of healthy adults walking a range of distances typical of gait tests (Fig. 2). Previously published data were re-analyzed in the context of gait speed, with methods briefly summarized here (for details see Carlisle and Kuo, 2023).

**Fig. 2.**
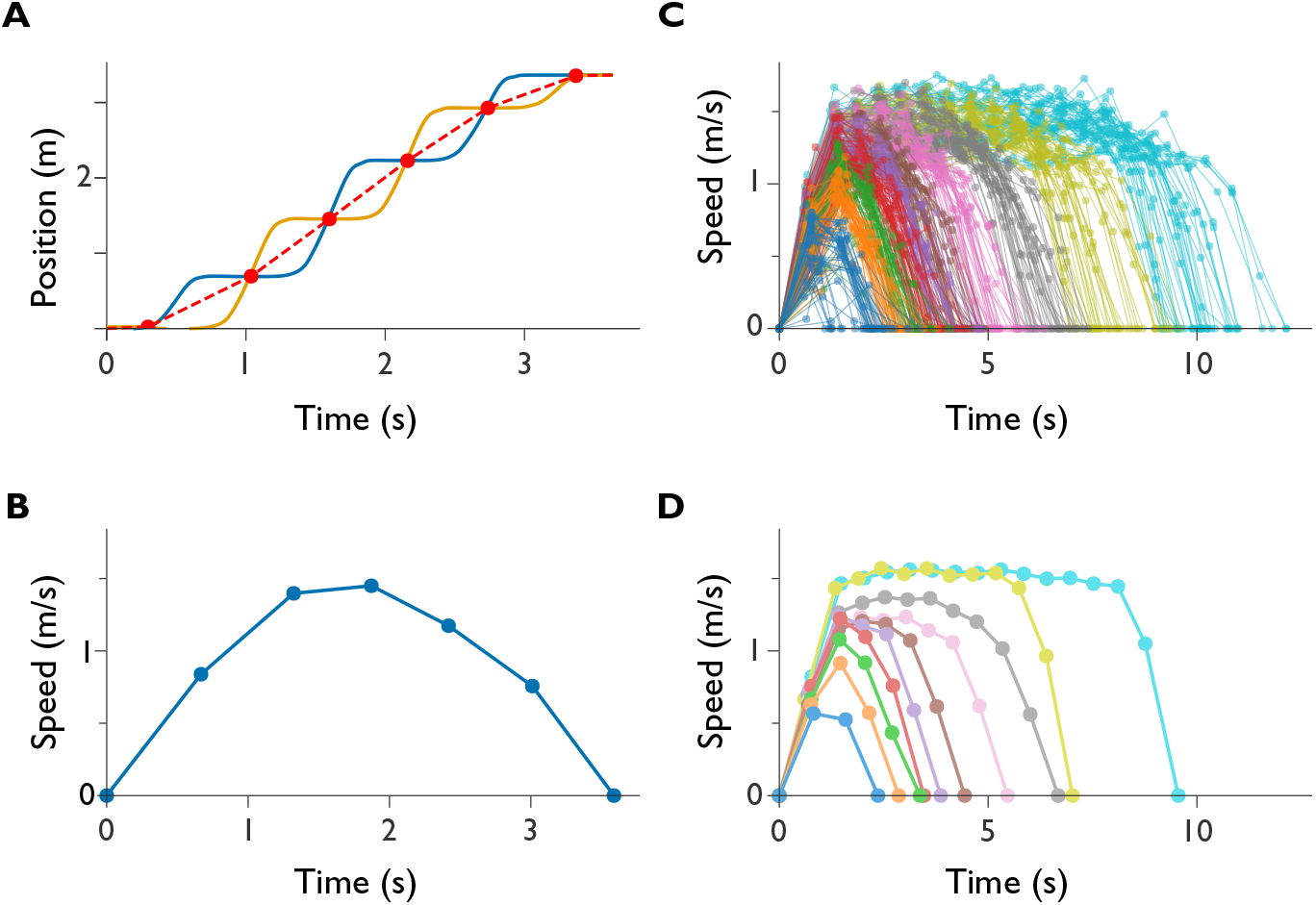
Observations of short walking tests ranging 1 – 12 m (Carlisle & Kuo, 2023). (A) Forward position vs. time for both feet (one trace each). The average of the two trajectories (dashed line) represents whole-body forward position. (B) Step-to-step forward speed vs. time, computed as body displacement divided by step duration. Filled circles indicate individual step speeds (assigned to foot position intersection instances); lines connect steps within a trial. (C) Speed vs. time walked for ten healthy adults walking ten target distances (color-coded). Each trace represents one of four trials per distance (400 total). (D) Mean speed profiles for each distance, plotted for illustration only and not used in analysis.

Subjects (N = 10, 6 male and 4 female, 24 – 38 yr) performed self-selected walking bouts on level ground at ten total target distances (about 1.1 m – 12.7 m, four trials each) marked by approximate targets on the ground, with static start and stop at upright standing. The forward foot positions (Fig. 2A) were measured with foot-worn inertial measurement units (IMUs) using inertial navigation procedures to reduce integration drift (Rebula et al., 2013). A speed was computed per step, defined from times when one foot passed another (Fig. 2A) as the distance divided by time interval between crossings, yielding a trajectory of speed vs. time for each trial (Fig. 2C). To visualize typical behavior, average trajectories were computed for each target walking distance (Fig. 2D), but all other analyses used individual trials. Each trajectory yielded total distance walked *D*, duration *T*, and peak speed *V* for each trial.

We sought compact empirical functions to describe how peak speed and total duration vary with total walking distance. The goal was to determine whether a single “steady” speed could explain performance across all distances, and whether that speed could serve as an objective measure independent of test procedure. This was approached with two ad hoc curve fits intended to summarize all of each subject’s trials.

The first curve fit was to represent how an individual’s peak speed increases with and saturates with total distance (Carlisle and Kuo, 2023),

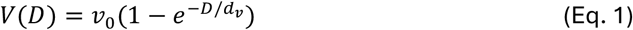

where *v*_0_ is the saturating speed and *d*_*v*_ a distance constant. We tested whether this saturating exponential described all walks, including steady ones for longer distances and slower and unsteady speeds for shorter distances. This would support the definition of an individual’s *preferred speed* as the *v*_0_ identified from all of their trials (40 per subject) across all distances. The complementary distance constant *d*_*v*_ quantifies how peak speed is approached over shorter distances, and thus describes how the individual accelerates. It takes about 2.3*d*_*v*_ to reach 90% of saturating speed.

The second fit was to represent how total duration approaches a linear relationship with total distance,

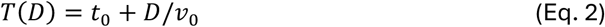

where *v*_0_ is the same saturating speed above and parameter *t*_0_ describes the total time spent accelerating and decelerating. We discarded a previous, nonlinear saturation term for very short distances (Carlisle and Kuo, 2023), which was found difficult to fit precisely to individual data, and to have negligible effect on *v*_0_ when applied to walking distances of at least 2.2 m (at least 26 trials per subject). We tested whether an individual-specific *v*_0_ could be identified independently by the two curve fits (Eqs. 1 & 2), which would indicate an objective quantity.

In addition, we also tested whether *v*_0_ could be identified from a small number (*n* = 3 to 5) of trials commensurate with most test conditions. We used a 4-iteration repeated hold-out procedure, where each iteration used one random trial from each of three target distances (3.8 m, 5.1, 7.0 m), until all four trials has been sampled once (without replacement), yielding four independent estimates for cross-validation. This “few-distance” procedure was also conducted for *n* = 4 and 5 targets (adding distances 9.1 m and 12.7 m).

To relate fitting parameters to current gait measures, we emulated conventional test protocols using the same dataset. We used common timing distances of 4 m and 10 m with **static starts**, applied to trials of total distance 4.5 m to 5.5 m (39 trials) including finish line (Bohannon and Wang, 2019) and greater than 10 m (44 trials), respectively. We also used all sufficiently long trials in **dynamic tests**, with both dynamic start and stop distances of both 1 m and 2 m. Gait speeds were calculated as timing distance divided by duration. Based on observed speed trajectories (Fig. 2D), we expected all gait tests to be correlated with saturating speed, albeit with tendency to underestimate it, and less so with longer dynamic start and stop.

We compared these protocol-based speeds to the saturating speed *v*_0_ from model fits. Bias and variance were computed to quantify the difference between conventional test estimates and *v*_0_, defined as mean and standard deviation phases based on first and last crossing of instantaneous speed with 90% of saturating speed (0.9*v*_0_), a practical threshold allowing for noisy fluctuations in seemingly steady walking. Bias and variance were similarly computed for the few-distance fits with hold-out cross-validation.

## Results

The empirical curve fits describe speeds and durations well across all distances. We first summarize the trajectory data and curve fits, and the justification for defining *preferred speed*. We next examine how short distances emphasize acceleration and deceleration, how conventional gait tests yield biased speed estimates, and how few trials are needed to determine preferred speed reliably.

### Peak speed and duration increase systematically with total walking distance

Subjects walked in a consistent manner across all distances (Fig. 3). The saturating speed fits (Fig. 3A) confirmed that speed trajectories (Fig. 2C) diverged from the conventional assumption of steady-state walking (Fig. 1B). For shorter distances, trajectories followed a rounded, inverted-U profile (Fig. 2D), with steady speed attained only momentarily or not at all. With increasing distance, peak speeds became both higher and more sustained, revealing a sustained steady interval lasting several steps for distances of at least 12 m— longer than most conventional gait tests. Walking duration (Fig. 3B) also increased systematically and approximately linear with longer distances.

**Fig. 3.**
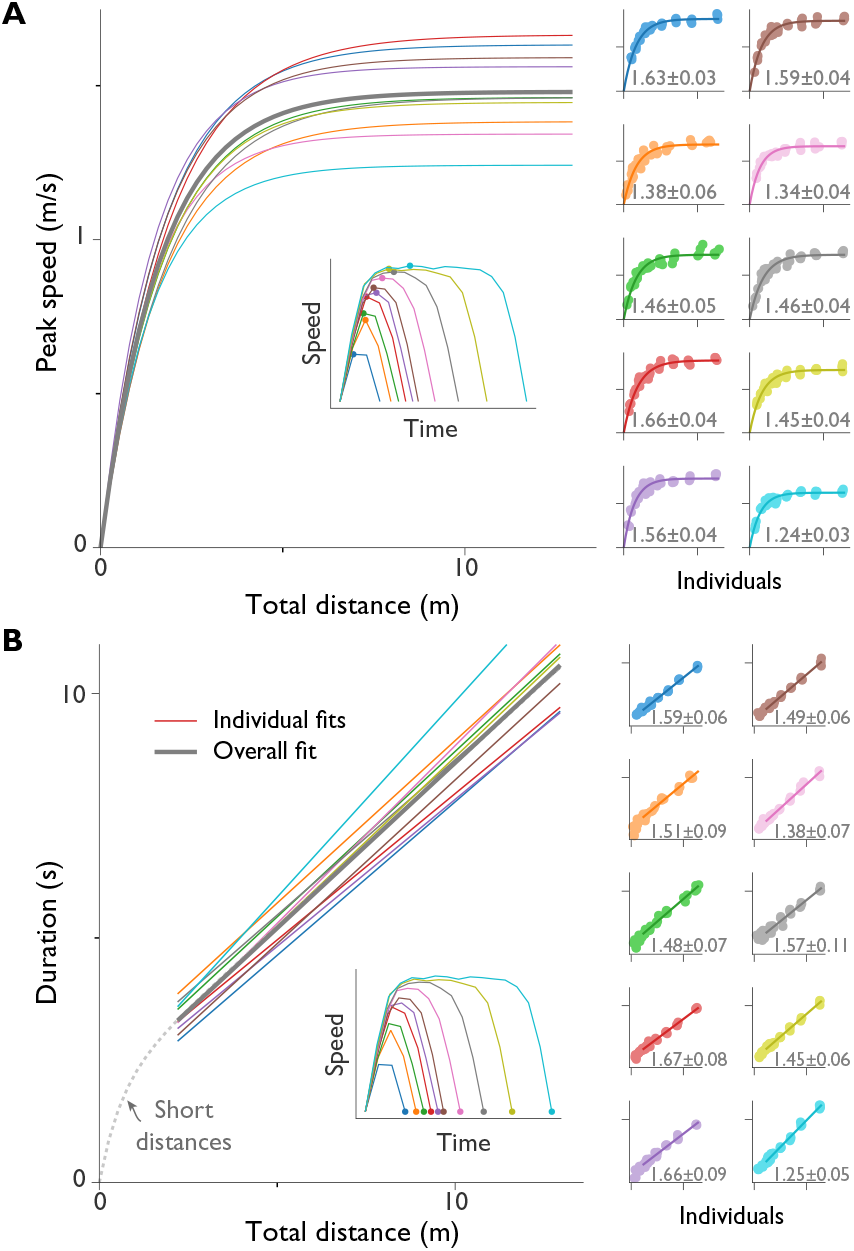
Peak speed and duration across walking distances, with corresponding curve fits. (A) Peak speed vs. total walking distance (left), fitted using a saturating exponential (Eq. 1; *R*^2^ = 0.94). Individual-subject fits (N=10) and data (ten distances × four trials) are shown in right panels. (B) Total duration vs. distance (left), fitted with linear relationship (Eq. 2; *R*^2^ = 0.98). Individual fits and data are shown in right panels. Duration fits exclude short distances < 2.2 m where relationship is curved (dashed line). Both curve fits converge to a steady quantity termed *preferred speed:* the saturating asymptote of peak speed in (A) and the inverse slope of distance-duration in (B). Right panels show each individual’s preferred speed ± m.e. (margin of error, or half-width of 95% c.i.). Left insets illustrate the speed and duration measures used for fitting.

### Peak speed saturates exponentially at longer distances

Peak speed *V* followed an approximately exponential saturation with total distance *D* (Eq. 1), with individual fits yielding asymptotic steady-state speeds *v*_0_ of 1.24 – 1.66 m ⋅ s^−1^ across subjects (1.48 ± 0.13 m ⋅ s^−1^, mean ± s.d., N=10), agreeing well with data (*R*^2^ = 0.94 ± 0.02).

### Saturating speed characterizes an individual’s preferred speed

Although each short walking bout had a unique speed, the saturating speed was robust across fitting methods (Eqs. 1 and 2), measured quantities (*V* or *T*), and walking distances. It was also stable under cross-validation, with hold-out resampling yielding a low standard deviation (0.08 m ⋅ s^−1^ s.d. for *n* = 3).

The slope of the duration-distance relationship (Fig. 3B) yielded comparable asymptotic speeds of 1.25 – 1.67 m ⋅ s^−1^ (1.51 ± 0.13 m ⋅ s^−1^; Eq. 2, *R*^2^= 0.98 ± 0.01). The saturating speed and linear duration fits also agreed with one another to within about 2%, with intraclass correlation 0.86. The saturating exponential yielded narrower confidence intervals (maximum margin of error m.e. 0.06 m ⋅ s^−1^ and 0.11 m ⋅ s^−1^, respectively; m.e. is half-width of 95% confidence interval for each subject’s fit), indicating higher precision. We therefore used the saturating exponential *v*_0_ as reference for bias and variance computations.

These results show that *v*_0_ characterizes an individual rather than a protocol, and can therefore serve as an objective, reproducible descriptor of preferred walking speed.

### Distance constant characterizes approach to preferred speed

The fitted constant *d*_*v*_ describes the shape of the saturation curve (Fig. 3A). Fitting individual data to Eq. 1 yielded *d*_*v*_ ranging 1.40 – 1.86 m (about ±14.4%) across subjects (1.63 ± 0.16 m, mean ± s.d.). This was equivalent to a range of 3.2 – 4.3 m total distance for subjects to reach 90% of preferred speed, and 4.2 – 5.6 m for 95%. Although preferred speed *v*_0_ spanned a similar ±14.3% range, the two were only loosely correlated, *ρ* = 0.55. The saturation curve’s amplitude (peak speed) was therefore only partially predictive of its shape (distance constant), and both parameters are needed to characterize individuals well.

### Acceleration and deceleration dominate shorter walks

Most of a shorter walking trial was occupied by acceleration and deceleration (Fig. 4A), with only about 67% of trials up to 5 m attained the threshold of 90% saturating speed (Fig. 4B), generally with a momentary peak (Fig. 2D). Across all distances, deceleration comprised approximately the same proportion as acceleration. The remaining proportion, for walking above threshold, only occupied about half the time of trials of at least 7.5 m, and only about 60% for 12 m walks.

**Fig. 4.**
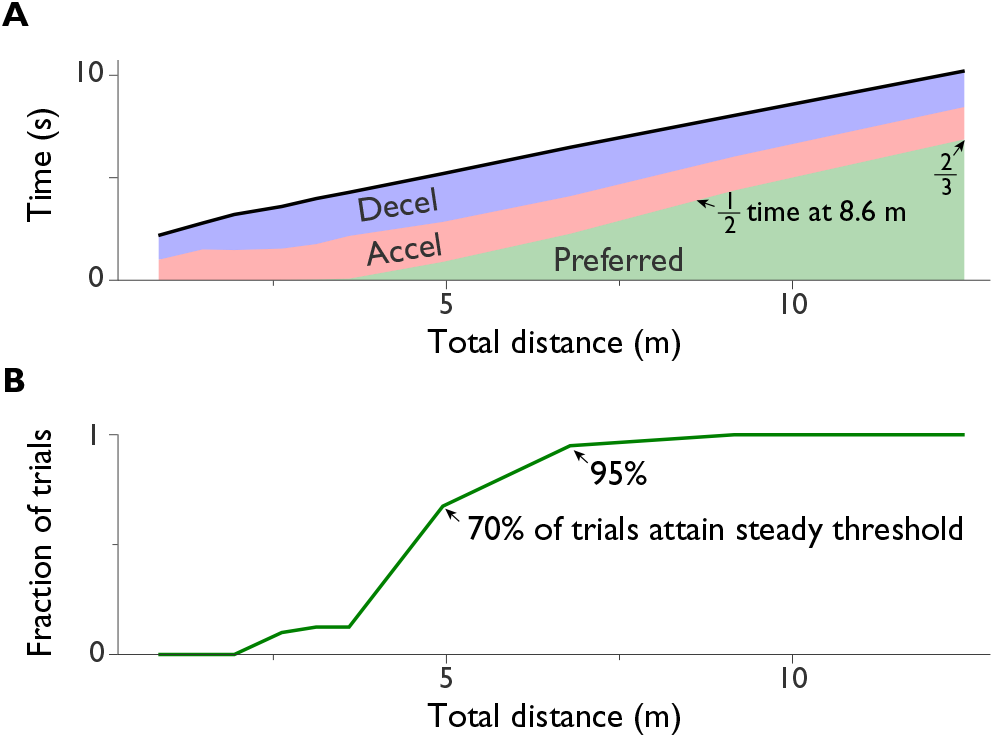
Time spent in acceleration, preferred steady speed, and deceleration phases as a function of total walking distance. (A) Duration of each phase (color-coded regions). Threshold for preferred steady walking defined as 90% of preferred speed (0.9*v*_0_). Walking at preferred speed occupies about half of and 8.6 m walk, and about two-thirds of a 12 m walk. (B) Fraction of trials reaching preferred speed threshold vs. total walking distance. About 70% of 5 m walks reached threshold, and 95% of 6.8 m walks. Data are from healthy adults (N = 10) walking ten target distances, four trials per distance.

### Short conventional gait tests underestimate steady speed

Application of conventional gait test measures to these data showed a negative bias relative to saturating speed (Fig. 5A). Static starts yielded the greatest underestimation, averaging -0.36 ± 0.08 m ⋅ s^−1^bias for 4 m tests and -0.14 ± 0.05 m ⋅ s^−1^ for 10 m (mean ± s.d. of residuals from *v*_0_; 39 and 44 trials, respectively).

**Fig. 5.**
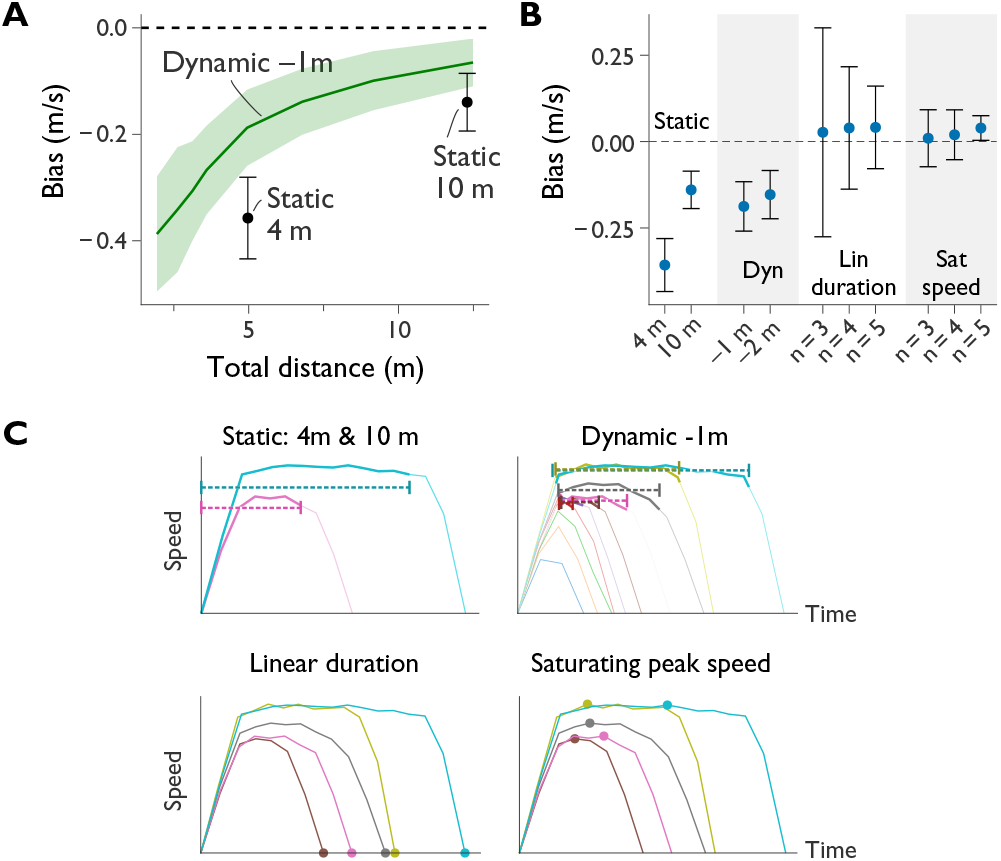
How test protocol affects saturating speed estimate *v*_0_, expressed as a bias relative to preferred speed. (**A**) Bias for static-start and dynamic-start/stop protocols plotted against total walking distance. “Static” 4m and 10m refer to timed distances of standard gait tests starting from rest. “Dynamic –1 m” indicates tests with 1 m start and stop intervals excluded from timing. Error bars and shaded area show standard deviation of residuals across cross-validation trials and subjects. (**B**) Bias for all metrics, also including fits for linear-duration and saturating peak speed. Fits were performed with n=3, 4, or 5 different total distances. Error bars show standard deviation of cross-validation residuals. (**C**) Representative speed-time trajectories illustrating timing intervals and resulting average speeds (dashed lines) for each method. Thick solid lines emphasize the data used for static and dynamic protocols. For linear duration and saturating speed fits, filled symbols indicate the duration and speed data.

Dynamic protocols yielded decreasing bias with distance (Fig. 5A). At a distance of 6.8 m, a 1m dynamic start and stop yielded bias -0.14 ± 0.06 m ⋅ s^−1^, comparable to a 10 m static start. A 2 m dynamic start/stop yielded only slightly less bias (Fig. 5B). Both static and dynamic protocols yielded comparable variance, decreasing with total distance.

### Preferred speed can be estimated with few walking trials

Fitting the saturation curve to only three trials of differing distances (3.8 m, 5.1 m, 7 m) yielded low bias (0.039 ± 0.036 m ⋅ s^−1^) and comparable bias relative to conventional gait tests (Fig. 5B). Adding a fourth and fifth distance (9.1 m and 12.7 m, respectively) further reduced variance. The linear duration fits yielded similar bias but greater variance. (Variances computed with 4-fold cross-validation per subject.)

## Discussion

Walking speed rarely remains steady over short distances. Instead of a constant plateau vs. time or distance as often assumed (Fig. 1B), the speed trajectory rises to a momentary peak and then immediately falls (Fig.2). The height of that peak increases with total distance, and only plateaus at a consistent level for distances longer than most gait tests (Fig. 3). This has implications for how gait tests should be interpreted and steady speed should be defined, and yields new insights on improved measures of walking.

These findings show that gait speed is more elusive than previously recognized. Nearly all of the walking time for shorter distances is spent accelerating and decelerating, and many trials never reach an eventual steady speed (Fig. 4). Even with dynamic starts and stops, the measured average speed generally remains biased below the individual’s characteristic speed (preferred speed, Fig. 5). In practical terms, gait tests shorter than about 10 m primarily reflect how quickly a person can speed up and slow down. Longer tests do increasingly reflect a steadier characteristic speed, but in a complex interplay with acceleration. This may help explain why measured gait speeds can vary across test protocols, making them difficult to interpret and limiting their objectivity.

To address this limitation, we defined preferred speed as the saturation limit of the peak speed-distance relationship. It readily integrates information across multiple test distances, yielding a measure independent of the specific protocol and distances used. Preferred speed is also independent of estimation method, because it can be estimated from independent distance-duration fits with low cross-validation error (Fig. 5). In practice, it can be estimated from a few walking trials at different distances, or directly from a single longer dynamic test (albeit with greater bias, Fig. 5). This measure therefore provides an objective and method-independent summary of an individual’s characteristic steady speed.

Beyond preferred speed, individuals also differ in how they accelerate. This additional dimension was quantified by distance constant *d*_*v*_ (Eq. 1), where greater values mean longer distance to attain preferred speed, and thus slower acceleration. The loose correlation with speed is exemplified by a few individuals who had faster preferred speeds than others yet accelerated more slowly (see crossings of individual curves in Fig. 3). This quantifies a long-recognized but previously unmeasured distinction: short gait tests with static starts emphasize acceleration, whereas longer dynamic ones emphasize steady-state speed. The interpretation of gait test outcomes might be improved by quantitatively distinguishing between acceleration and sustained walking performance.

These findings suggest several refinements for conventional gait tests. First, both start and stop procedures should be clearly specified and reported, since they are of equal importance. Instructions such as “walk past the finish line” invite interpretation, unlike a clear stopping point. Second, it should be recognized that short gait tests usually do not attain steadiness. Although dynamic start and stop conditions can mitigate this, steadiness should only be claimed when supported by quantitative evidence. Steadiness is also not critical to a gait test’s clinical importance. Third, when additional trials are feasible, varying the total distance can yield more information than repeating a single one, for example facilitating curve fit analysis as shown here. Finally, the timing interval could be considered incidental to the more ecological task of walking between defined start and stop locations. A concise example incorporating these principles might read: “Subjects were instructed to walk three total distances of 5 m, 9 m, and 11 m, each marked by stop points. They were timed from a static start until crossing a line 1 m before the stop point.”

Better quantification could also be incorporated into gait tests. Automatic measurements no longer require specialized equipment such as gait mats or motion capture, and can now be performed by IMUs or portable digital video cameras at modest cost ($100 USD) and minimal set-up or training time. This could not only improve timing precision for conventional tests but also record position or speed trajectories (e.g., Fig. 2), which could verify when steady speed is attained and support estimation of preferred speeds and distance constants. Richer trajectory information can thus be obtained with only slight alterations to existing test procedures.

This study has several limitations. We measured gait in only a modest number of healthy subjects, and did not include important populations such as older adults or those with infirmities. As such, effects such as distance-dependent speed saturation have not been proven for many conditions. We expect some populations to attain steady speed differently, and perhaps more slowly, than reported here. We also tested only a limited range of distances here, and note the need for more data for longer distances such as 20 m or 100 m. Indeed, steadiness itself appears variable over yet longer distances (Lindemann et al., 2021). We also conducted point-to-point walking tasks rather than following the literal instructions of conventional gait tests. We consider the results to be indicative of such tests but not exactly equivalent. We also anticipate that there are other dimensions to human gait in addition to the preferred speed and distance constant examined here. This study should therefore be regarded as an implication of better ways to characterize gait speed, that merit more testing on more populations.

These results do not diminish the value of conventional gait tests, which remain useful clinical indicators correlated with health outcomes. Rather, we hope to clarify what such tests actually measure under different conditions, and suggest a framework for making them more quantitative, objective, and interpretable, akin to established vital signs.

Recognizing the two complementary dimensions—preferred steady-state speed and acceleration capacity—may allow future assessments to disentangle the multiple physiological and biomechanical factors that together determine walking performance.

